# Representational geometries reveal differential effects of response correlations on population codes in neurophysiology and functional magnetic resonance imaging

**DOI:** 10.1101/2022.11.17.516856

**Authors:** Zi-Jian Cheng, Wen-Hao Zhang, Ru-Yuan Zhang

## Abstract

Two sensory neurons usually display trial-by-trial response correlations given the repeated representations of an identical stimulus. The effects of such response correlations on population-level sensory coding have been the focal contention in computational neuroscience over the past few years. In the meantime, multivariate pattern analysis (MVPA) has been the leading analysis approach in functional magnetic resonance imaging (fMRI), but the effects of response correlations in voxel populations remain underexplored. Here, instead of conventional MVPA analysis, we calculate linear Fisher information of population responses in human visual cortex and hypothetically remove response correlations between voxels. We found that voxelwise response correlations generally enhance stimulus information, a result standing in stark contrast to the detrimental effects of response correlations reported in neurophysiological literature. By voxel-encoding modeling, we further show that these two seemingly opposite effects actually can coexist. Furthermore, we use principal component analysis to decompose stimulus information in population responses onto different principal dimensions in a high representational space. Interestingly, response correlations simultaneously reduce and enhance information on high- and low-variance principal dimensions, respectively. The relative strength of the two antagonistic effects within the same computational framework produces the apparent discrepancy in the effect of response correlations in neuronal and voxel populations. Our results suggest that multivariate fMRI data contain rich statistical structures that are directly related to sensory information representation, and the general computational framework to analyze neuronal and voxel population responses can be applied in many types of neural measurements.

## INTRODUCTION

A fundamental challenge in systems neuroscience is to understand how sensory information (e.g., the orientation of a line segment) is encoded by evoked neural responses. It has been well-established that even a simple sensory stimulus can evoke the responses of a large group of neurons, a phenomenon termed *population codes* in computational neuroscience1,2. Previous theoretical and empirical neurophysiological studies have identified several key aspects of population codes with respect to sensory coding, such as tuning functions and response variability of individual neurons, and trial-by-trial response correlations between neurons 3. Understanding the relationship between these response properties and the fidelity of sensory coding is vital to uncover the neural mechanisms of bottom-up and top-down sensory processing.

The most distinguishable feature of population codes, as compared to single-unit recording data, is the trial-by-trial response correlations (RC) between units. How correlated variability affects the fidelity of population codes has been a focal contention in neuroscience in the past few years. On one hand, theoretical studies have shown that the effects of neuronal RCs depend on many factors, such as the form of RC, tuning heterogeneity, or its relevance to behavior 1,2,4-7. On the other hand, empirical physiological studies have consistently shown that improved behavioral performance is accompanied by a reduction of RC in an array of cognitive processes, such as locomotion 8, task engagement 9, wakefulness 10, and perceptual training 11, implying the detrimental role of RC in sensory coding. RCs between neurons have also been shown pivotal in top-down attentional modulations 12,13.

Unlike invasive neurophysiology, functional magnetic resonance imaging (fMRI) is a noninvasive technique and naturally measures the responses of many units in the brain, albeit the basic unit is voxel, not neuron. Both fMRI and neurophysiology research thus face the same challenge: how can we elucidate the mechanisms of sensory processing from high-dimensional data? Henceforth, we use “population codes” as a unified term for both neurophysiology (i.e., a population of neurons) and fMRI (i.e., a population of voxels). fMRI research fully utilizes the advantage of multivariate responses, as evidenced by the fact that multivariate pattern analysis (MVPA) has become a mainstream approach to decipher sensory information 14. In a standard MVPA (e.g., orientation decoding), researchers first estimate single-trial population activity (i.e., beta weights from general linear modeling in fMRI) towards each stimulus (similar to measurements of neuronal firing rate in animal physiology). In conventional fMRI literature, correlations between the trial-by-trial responses (i.e., beta weights) of two voxels (Fig. 1D) are called *Beta-series correlation* 15,16. We particularly emphasize that Beta-series correlation is neither resting-state functional connectivity calculated by correlating the time series of two voxels during rest 17, nor task-based functional connectivity (or background functional connectivity documented in some literature) calculated by correlating the residual time series from general linear modeling 18. Similarly, trial-by-trial spike-count correlations between neurons are termed “noise correlation” in neurophysiology 19. Though given different names, both Beta-series correlation in fMRI and noise correlation in neurophysiology describe the trial-by-trial response correlations between two units, and we henceforth unify them as *response correlation* in this paper. In fMRI, the quality of stimulus encoding is usually operationally defined as multivariate decoding accuracy—that is, a higher decoding accuracy indicates higher fidelity of stimulus coding 20,21. This multivariate decoding approach has been widely used to study the neural mechanisms of a variety of cognitive processes 14,22. For example, attention and learning can improve the decoding accuracy of some basic visual features (e.g., orientation, color, or motion direction) in the human visual pathway 23. Compared to rich theoretical and empirical evidence for the effects of RC in neurophysiology, little is known with respect to how voxelwise RCs impact MVPA decoding accuracy in fMRI. Unveiling the effects of RCs directly speaks to the mechanisms of many cognitive processes. For example, if decoding accuracy is elevated by a particular cognitive process (e.g., attention), it is unclear whether it warps response correlations (Fig. 2C) or simply reduces the response variability of single voxels (Fig. 2D).

**Figure 1.**
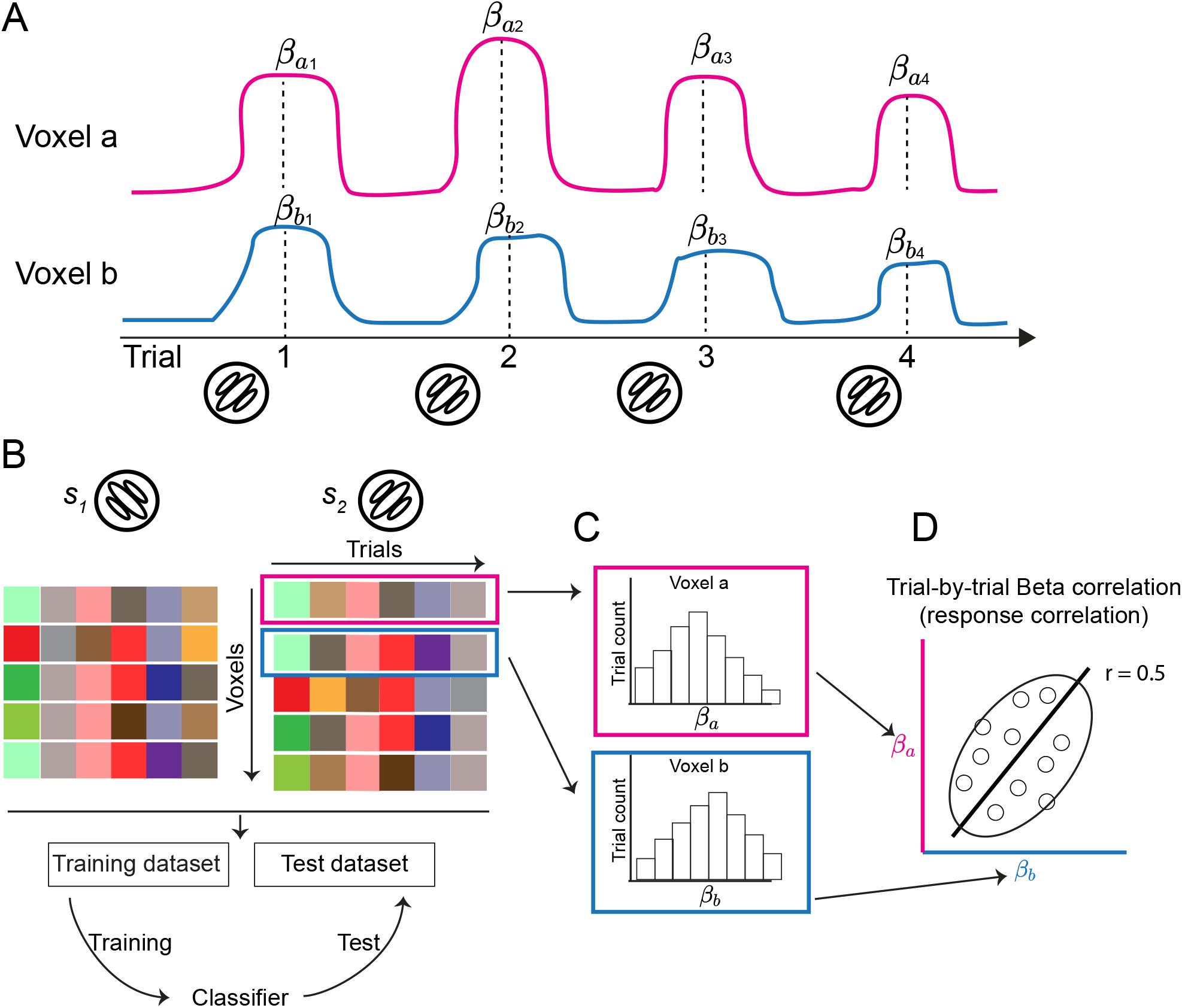
Trial-by-trial response correlation in multivariate fMRI data. ***A***. Estimations of single-trial voxel responses. Margenta and blue lines represent the time series of voxels *a* and *b* across four trials of the same grating stimulus. For MVPA, the stimulus in each trial is modeled as a single predictor in general linear modeling to estimate single-trial voxel activity (i.e., beta weight). *β*_*ai*_ (*β*_*bi*_) indicates the response of the voxel *a* (respectively, voxel *b*) in the *i*-th trial. ***B***. In the problem of binary classification, multivariate voxel responses for each stimulus can be summarized as a voxel-by-trial matrix. Each item in this matrix is the activity of the *i*-th (row) voxel in the *j*-th trial (column). Combining the two data matrices for the two stimuli, a classifier can be trained on a training dataset and then evaluated on a test dataset. ***C***. A single voxel’s responses across trials (i.e., a row in a data matrix in ***Panel B***) is a random variable following some distribution. ***D***. Beta-series correlation can be visualized by the scatter plot of the responses of voxels *a* and *b* (i.e., two rows in a data matrix in ***Panel B***). The solid line is the correlation line, the empty dots represent the activity of individual trials. The ellipse illustrates the shape of the 2d response distribution, and the direction of the ellipse depicts a positive response correlation (e.g., r = 0.5 in this example) between the two voxels. We particularly emphasize that between-voxel Beta-series correlation is neither resting-state functional connectivity nor task-based functional connectivity calculated by correlating the residual time series after regressing out stimulus-evoked responses from general linear modeling.

**Figure 2.**
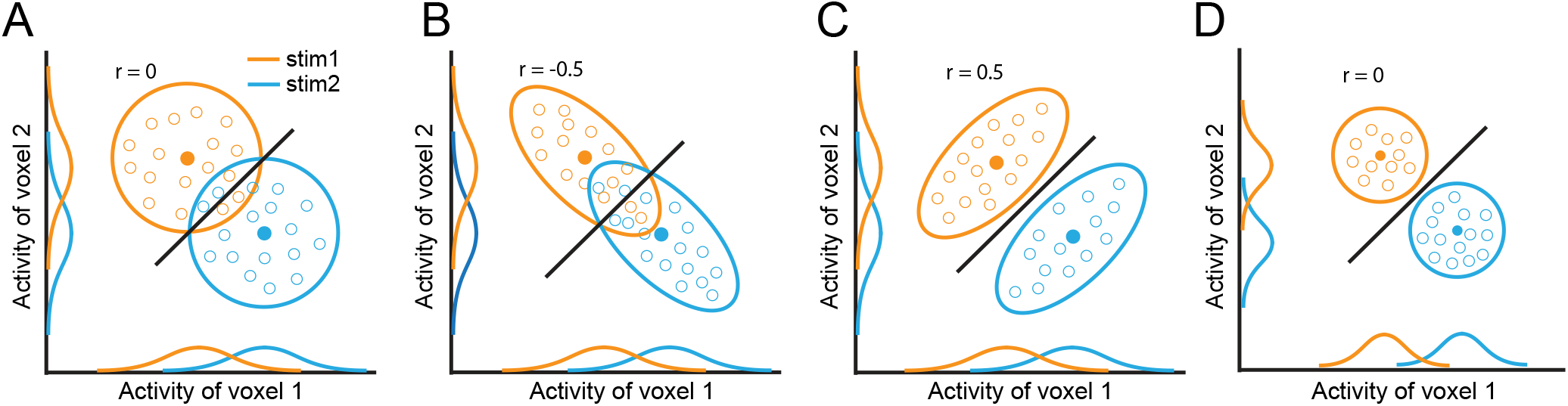
A two-voxel scenario showing the effects of RCs on decoding. The blue and orange circles or ellipses represent 2-d response distributions towards two stimuli. The solid dots represent the mean of the distributions, and the empty dots represent individual trials consisting of the distributions. The solid lines are classifiers. The 1-d distributions are the projection of the 2d distributions on the x- and y-axes. In this example, compared to the case of no RCs (e.g., r = 0, ***Panel A***), negative RCs impair (e.g., r = -0.5, ***Panel B***) and positive RCs improve (e.g., r = 0.5, ***Panel C***) decoding accuracy. Note that the example correlation values correspond to the 2-d distributions, not the classifier lines. The amount of stimulus information is directly linked to decoding accuracy. As such, stimulus information in ***Panels C&D*** is higher than in ***Panels A&B***. Importantly, the marginal distributions in ***Panels A-C*** are identical but the decoding accuracies are markedly different across the three cases. In ***Panel D***, the variance of the two distributions shrinks and consequently induces higher decoding accuracy. ***Panels C&D*** demonstrate that identical decoding accuracy can correspond to two distinct underlying representational geometries.

Here, we employ an information-theoretic approach instead of the conventional decoding approach to analyze the fidelity of stimulus encoding in multivariate fMRI data. In the case of classification, *stimulus information* here refers to the discriminability of two stimuli based upon trial-by-trial population responses. Specifically, we directly calculate linear Fisher information of stimuli—a standard metric to assess stimulus information in computational neuroscience— encoded by trial-by-trial multivariate voxel responses. Surprisingly, in contrast to the detrimental effects documented in neurophysiological literature, we find that voxelwise RCs primarily improve stimulus information in the human early visual cortex, and stimulus information follows a U-shaped function of RC strength. Although the effects or RCs appear drastically different in neuronal and voxel populations, we used a voxel encoding model to show that these two effects can coexist in visual systems, and we observed them just because brain signals are acquired at two different spatial scales (i.e., microscopic neurons and macroscopic voxels). We further analyze stimulus information in eigendimensions instead of original voxel dimensions and find that the apparent difference in the effects of RCs in neuronal and voxel populations share some common mechanisms: RCs reduce, and in the meantime, enhance information on high- and low-variance eigendimensions, respectively. These two antagonistic effects jointly determine the total stimulus information in a population and produce the U-shaped function observed in fMRI data. This unified mechanism of RC not only helps resolve the long-term debate on the effects of RC on sensory coding in neurophysiology, but also extends the conventional understanding of the computational nature of MVPA. Moreover, the methods to analyze population responses here are general and can be applied to a wide range of regimes for deciphering multivariate neural data.

## RESULTS

### Voxelwise response correlations enhance stimulus information in human early visual cortex

We analyzed the trial-by-trial voxel activity in early visual cortex when six human participants viewed gratings with four orientations (i.e., 0°, 45°, 90°, and 135°, see Methods and Fig. 3A). Like classical decoding experiments 21, each orientation was presented for several trials, allowing us to estimate single-trial activity of individual voxels. Instead of the conventional decoding approach, we calculated linear Fisher information—how well two stimuli can be discriminated based upon population responses—in early visual areas (i.e., V1-V3) of all subjects. The calculation of linear Fisher information requires estimations of the response mean and variance of individual voxels as well as the response correlation matrices between voxels (see Eqs. 1-3 in Methods). As we will show later, this information-theoretic approach is advantageous as it allows one to symmetrically manipulate RC strength in data and examine its consequences.

**Figure 3.**
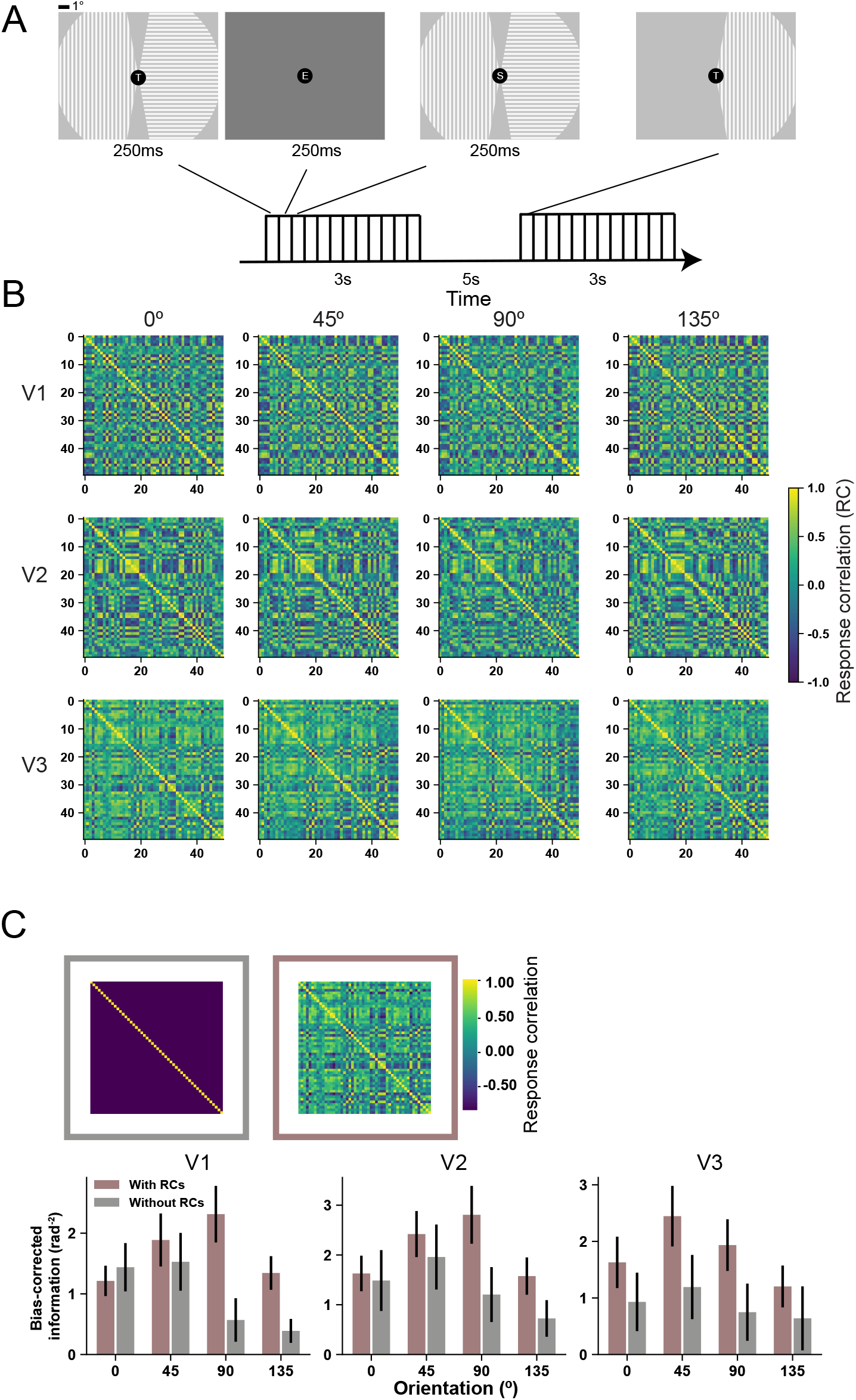
Voxelwise response correlations enhance stimulus information. ***A***. Schematic illustration of example trials in the orientation experiment. Two gratings are presented in each visual field. Their orientations are independently selected from four orientations (0º/45º/90º/135º). In some baseline trials (second trials shown above), only one grating is presented. Subjects are asked to attend and understand the letter stream presented at the fixation point. ***B***. example correlation matrices of V1-V3 in one hemisphere of one subject. These results show that the correlations between the same pool of voxels but across different stimuli are similar. In other words, the noise correlations between voxels are mostly stimulus invariant. ***C***. The left matrix is the identity matrix. The right matrix is an example response correlation matrix estimated from data. Hypothetical removal of response correlations between voxels (gray bars) reduces the amount of information as compared to the case that all response correlations are preserved (puce bars). The error bars represent S.E.M across 12 independent samples (6 subjects x 2 hemispheres). Note that the information here is bias-corrected using Eqs. 4&5 (see Methods).

To demonstrate the effects of trial-by-trial RCs, we compared stimulus information in each brain region under two regimes. In one regime, linear Fisher information was directly calculated based on empirically measured voxel responses with all voxelwise RCs being completely preserved. In the other regime, linear Fisher information was calculated with all voxelwise RCs being “hypothetically” removed. To do so, we manually set the RC matrices to identity matrices (see Eqs. 1-3 in Methods). Notably, this procedure forces the RCs between voxels to be zero (i.e., removal of all RCs) without changing the marginal response distributions (i.e., mean and variance) of individual voxels. Intuitively, this is equivalent to warping the two response distributions in Fig. 2B&C (i.e., RCs are non-zero) to Fig. 2A (i.e., RCs are zero). This method has been used to illustrate the effects of RCs in prior studies 24,25. Note that the estimated statistical properties (e.g., mean, covariance) given a limited number of units and trials may lead to bias in the estimation of information. We therefore calculated bias-corrected linear Fisher information under these two regimes (see Eq. 12 in Methods and ref. 24).

We found that the information with RCs being preserved is significantly higher than that with RCs being hypothetically removed, indicating that voxelwise RCs in general enhance stimulus information. Our finding here stands in contrast to the findings in neurophysiology as the majority of animal studies demonstrated the detrimental effects of neuronal RCs. We will explain the discrepancy in later sections.

### Information as a U-shaped function of correlation strength in voxel populations

The above analyses demonstrate the potential beneficial effects of voxelwise RCs by contrasting two regimes—voxelwise RCs are completely preserved or removed. The result shows that voxelwise RCs ostensibly enhance stimulus information. However, according to our previous theoretical work 26, the relations of RCs and stimulus strength may not be monotonic. To gain a full picture of the possible effects of RCs, we further used a “titration” approach: we calculated stimulus information while progressively manipulating RC strength. Specifically, all off-diagonal items of empirically measured RC matrices were multiplied by an RC coefficient (i.e., *c*_*νxs*_). The RC matrices are identity matrices if the coefficient is set to 0, and preserved with no change if the coefficient is set to 1 (Fig. 4A). In other words, the two regimes in Fig. 3 can be viewed as the two special cases (i.e., *c*_*νxs*_ = 1/0) of the “titration” analyses here.

**Figure 4.**
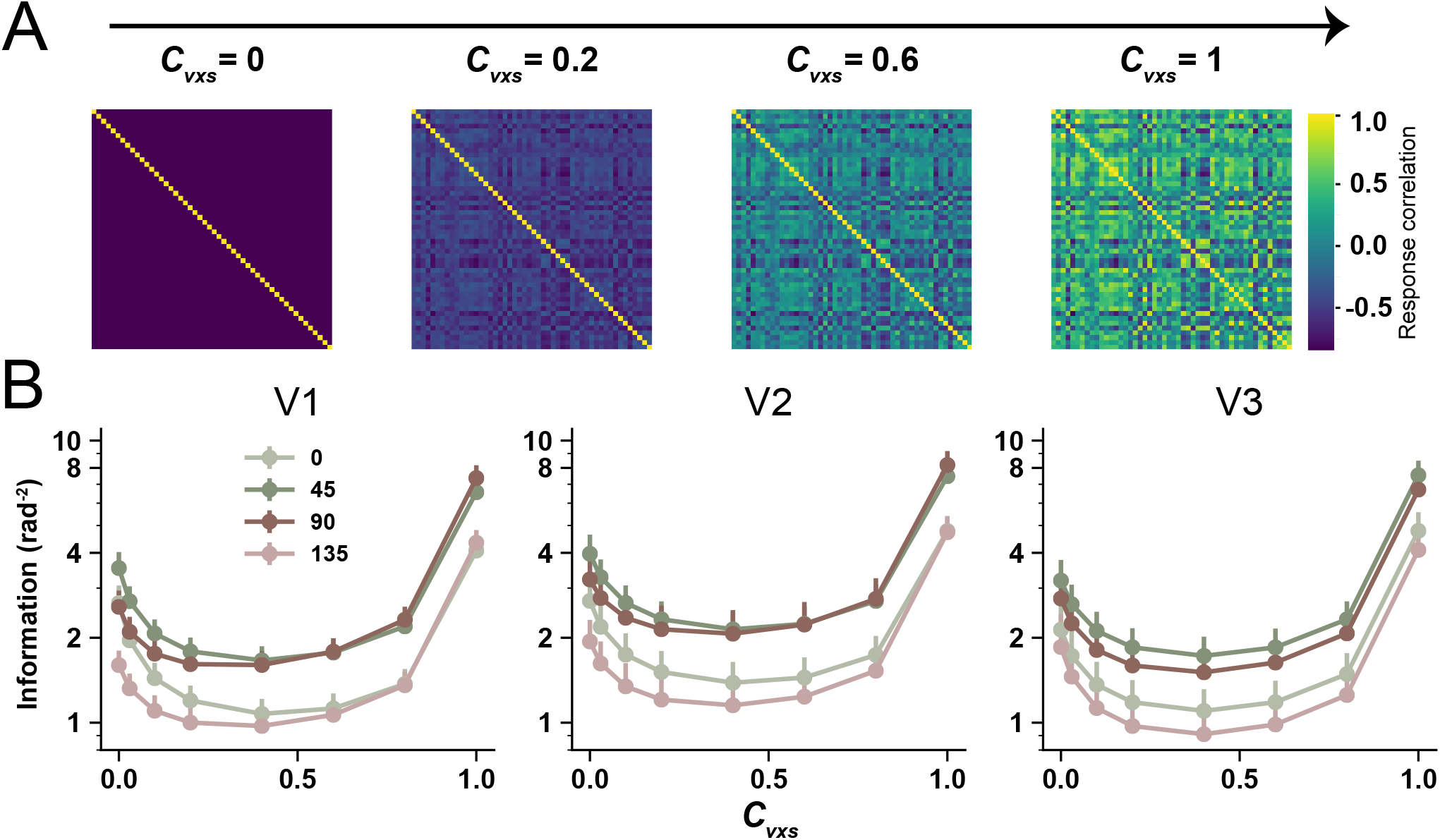
Stimulus information as U-shaped functions of voxelwise RC strength.*c*_*νxs*_ is the RC coefficient controlling RC strength. ***Panel A*** depicts that the off-diagonal items in RC matrices emerge as *c*_*νxs*_ increases. ***Panel B*** depicts information as U-shaped functions of *c*_*νxs*_. Note that the two conditions in Fig. 3 correspond to the first and the last data points of the curves shown here. But Fig. 1 shows bias-corrected linear Fisher information (Eq. 12), and here information is calculated using Eq. 1 without bias-correction because we are interested in the changing trend rather than the absolute amount of information. Error bars in ***Panel B*** are S.E.M across 12 independent samples (6 subjects x 2 hemispheres).

Interestingly, we found that although RCs improve information when all RCs are preserved, the effects of RCs are non-monotonic—stimulus information first declines then rises as increasing RC strength, manifesting as a U-shaped function. The ebb and flow can reach a half and double magnitude of that without RCs. This result held for all four orientations and in all three ROIs. Note that a few previous studies investigated the effects of RC by shuffling population responses across trials to remove RCs and then comparing decoding accuracy between unshuffled and shuffled data 12. That approach is similar to the analyses in Fig. 3. That shuffling approach, however, has two major shortcomings: 1) it has been proven to be less efficient when the number of trials and units are relatively small^24^, and 2) it cannot easily reveal the full changing trend of information when RC strength is progressively manipulated (e.g., the U-shaped function observed here).

### Voxel-encoding models explain the discrepancy between the effects of response correlations in neuronal and voxel populations

Why does stimulus information exhibit a U-shaped function as increasing RC strength? And more importantly, why do the effects of RCs manifest differently in neurophysiological and fMRI data? One notable difference between neuronal and voxel activity is that voxel activity is believed to reflect the aggregations of activity of many neurons. We next show that this neuron-to-voxel activity summation naturally produces the observed U-shaped function.

We first seek to replicate the classical detrimental effects of neuronal RCs using a neuron-encoding model and applied the same data analyses to fMRI data. One well-established finding in neurophysiology is that the response correlation between two neurons is proportional to their tuning similarity, called *tuning-compatible RC*2. In the neuron-encoding model, we simulated 50 orientation-selective neurons with Poisson response statistics, and assumed the tuning-compatible RCs between the neurons (Fig. 5A, see Eq. 6). We also simulated a control type of RCs, denoted as *tuning-irrelevant response correlations*. For tuning-irrelevant RC, the items in RC matrices are identical to those in tuning-compatible RC matrices but rearranged across columns and rows such that the RC between a pair of voxels bear no resemblance to their tuning similarity 27 We found that tuning-compatible RCs always impair information (green line in Fig. 5C) while tuning-irrelevant RCs always enhance information (orange line in Fig. 5C) in the neuron-encoding model. These results are consistent with a wide range of empirical or theoretical studies in neurophysiology 4,8,9,11.

**Figure 5.**
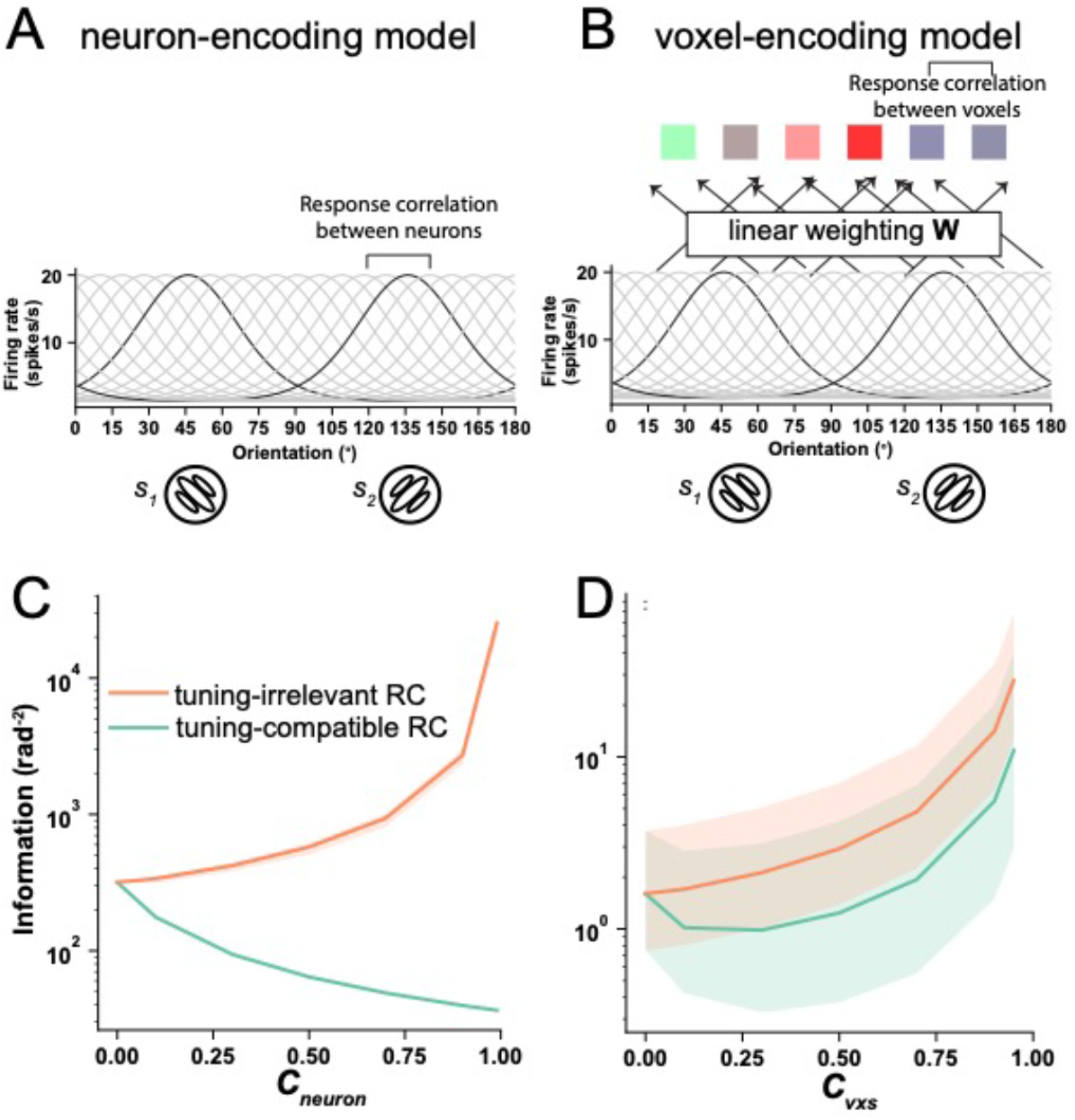
Neuron- (***Panel A***) and voxel-encoding (***Panel B***) models reproduce the effects of RCs reported in previous neurophysiological literature (***Panel C***) and our fMRI data (***Panel D***). *c*_*neuron*_ and *c*_*νxs*_ are the RC coefficients controlling the RC strength in the neuron- and the voxel-encoding model, respectively. The shaded areas of the pink curve in panel C and those of the two curves in panel D represent the 95% confidence interval for 100 simulations. Note that the green curve in ***Panel C*** (i.e., the neuron-encoding model with tuning-compatible RCs) has no variability because neuronal tuning curves and variance are fixed in this condition (see Methods for details).

We next turn to model the effects of voxelwise RCs on sensory information in fMRI data. Before doing so, we first investigated the structure of voxelwise RCs. It remains unclear whether tuning-compatible RCs also exist in fMRI data. To our best knowledge, only one study found evidence for the existence of tuning-compatible RCs in fMRI 27. We thus quantified the relationship between tuning similarity and response correlations in our empirical data (see Methods). We found that there is a significantly positive relationship between tuning similarity and response correlations in all 144 tests (all correlation p-values < 0.05, 6 subjects x 2 hemispheres x 3 ROIs x 4 stimuli, see details in Methods), substantiating the existence of tuning-compatible RCs in realistic fMRI data.

With tuning-compatible RCs being established in fMRI, we next established a voxel-encoding model that incorporates an additional layer of voxel activity on top of the neuron-encoding model and assumes voxel activity as the linear combinations of underlying neuronal activity (see details in Methods and Fig. 5B). This encoding-model approach has been widely used in fMRI studies on a variety of topics, including attention 28, memory 29, and learning 30. Note that, in this voxel-encoding model, the parameters (e.g., amplitude, baseline firing rate, tuning width) at the neuron-encoding stage are identical to those in the above neuron-encoding model. Similarly, we assumed that the voxelwise RCs 1) either are proportional to their tuning similarity (i.e., tuning-compatible RCs), or, as a control, 2) have no relationship with their tunings (i.e., tuning-irrelevant RCs). Interestingly, the model incorporating tuning-compatible RCs can well reproduce the U-shaped information function (the green line in Fig. 5D). Like neurons, the tuning-irrelevant RCs between voxels always improve information (the orange line in Fig. 5D). This quantitative voxel-encoding modeling suggests that the detrimental effects of RCs at the neuronal level and the potential beneficial effects of RCs at the voxel level can coexist, and we obtain opposite results just because brain activity is acquired at different spatial scales (i.e., neurons at the microscopic level and voxels at the mesoscopic level).

These results are also consistent with our previous theoretical work that voxel tuning heterogeneity contribute to the U-shaped function of the effects of voxelwise RCs 26. Unlike the relatively homogeneous tuning curves in the neuron-encoding model (i.e., similar peak, baseline of tuning functions) of single neurons, voxel tuning curves, as calculated by the aggregations of neuronal activity, are remarkably heterogenous (i.e., diverse baseline, response range). And such tuning heterogeneity attenuates the detrimental effects of RC. We will show this more formally in later sections.

Leveraging the voxel encoding model, we also investigated the nature of tuning-compatible RCs. Although we found tuning-compatible RCs in our empirical voxel responses, it remains elusive theoretically why they should exist at the first place. We hypothesized that the mapping from neuronal to voxel activity naturally produces tuning-compatible RCs among voxels. We first showed analytically that the weighting matrix **W** bridges the (co)variability from neurons to voxels (Eq. 13, also see 31). Then, using the voxel-encoding model above and a weighting matrix **W**, we calculated tuning similarity and response correlation between every pair of voxels in the voxel-encoding model, and examined their relations (see the example scatter plot in Fig. 6A). We repeated this calculation 1000 times and found a significant positive relation between tuning similarity and response correlations between voxels (bootstrap test: p < 0.013, Fig. 6B). In other words, we theoretically proved that there should exist tuning-compatible RCs between voxels as long as voxel activity is regarded as the aggregations of underlying neuronal activity.

**Figure 6.**
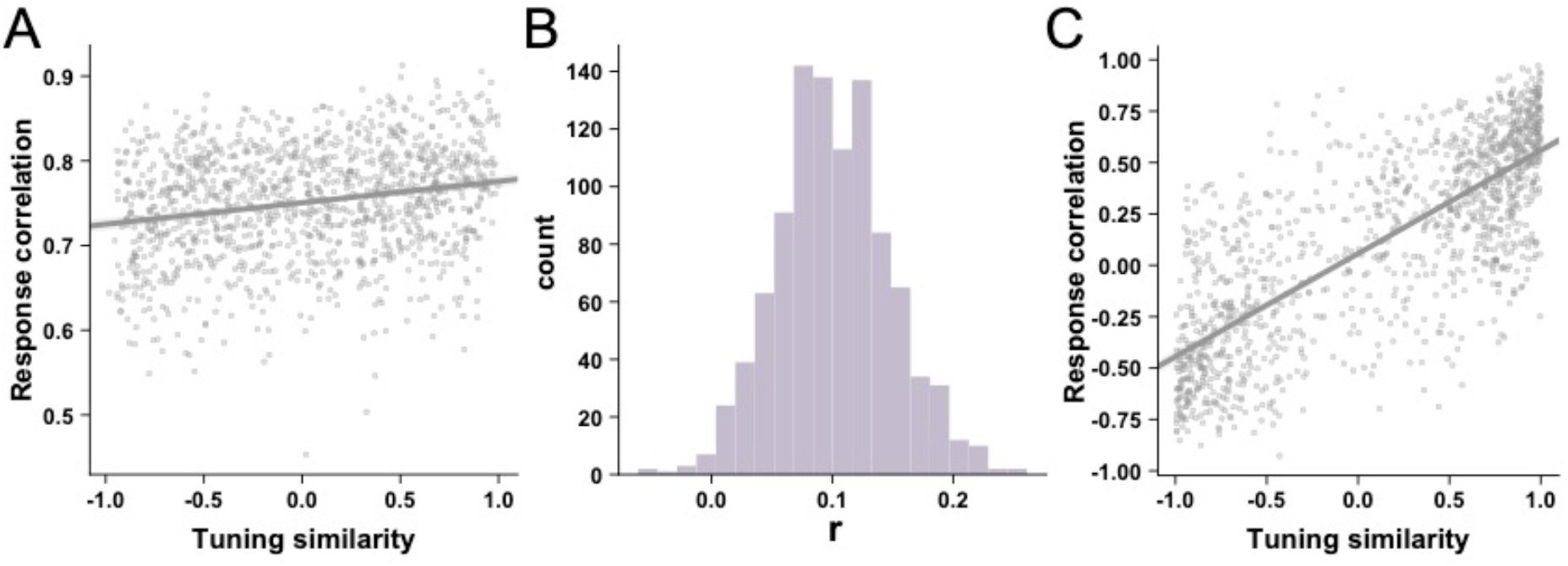
theoretical derivation and empirical measures of tuning-compatible RCs. ***A***. An example scatter plot of RCs (y-axis) and tuning similarity (x-axis) in one simulation of the voxel encoding model. ***B***. The distribution of the correlation value across 1000 simulations. In each simulation, we randomly generate a new weighting matrix **W. *C***. one example scatter plot of RCs (y-axis) and tuning similarity (x-axis) in a total of 144 cases (6 subjects x 2 hemispheres x 3 ROIs x 4 stimuli).

### Opposite effects of response correlations on stimulus information in neuronal and voxel populations revealed by eigen-information-decomposision analysis

The results above only demonstrate that neuron- and voxel-encoding models assuming voxel activity as aggregations of underlying neuronal activity can well-reproduce the effects of RCs observed in empirical data. However, what are the exact computational mechanisms underlying such discrepancy? Why does the transformation from the neuronal to the voxel level alter the effect of RC? To address this issue, we next introduce the method of information decomposition and then describe how it can serve as a unified mathematical framework to understand information coding in multivariate data.

The key formulation of information decomposition (as shown below see Eq. 1 and Eq. 12 in Methods) is to convert the calculation of linear Fisher information (*I*) in the original neuron/voxel space (*d***f** ^**T**^ * **Q**^−1^*d***f**) to the eigenspace 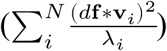:

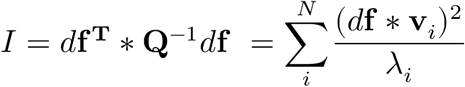

where *d***f** is the signal vector normalized to the absolute difference between the two stimuli (see Eqn. 2 and the red vector in Fig. 7C). **Q** is the averaged covariance matrices of voxel responses towards the two stimuli (see Eq. 3). **v**_*i*_ and λ_*i*_ are the *i*-th eigenvector and its corresponding eigenvalue (i.e., variance) of **Q**, respectively. This approach avoids the burdensome matrix inversion (**Q**^−1^) and represents the information of the whole population as the summation of the information along each eigendimension.

**Figure 7.**
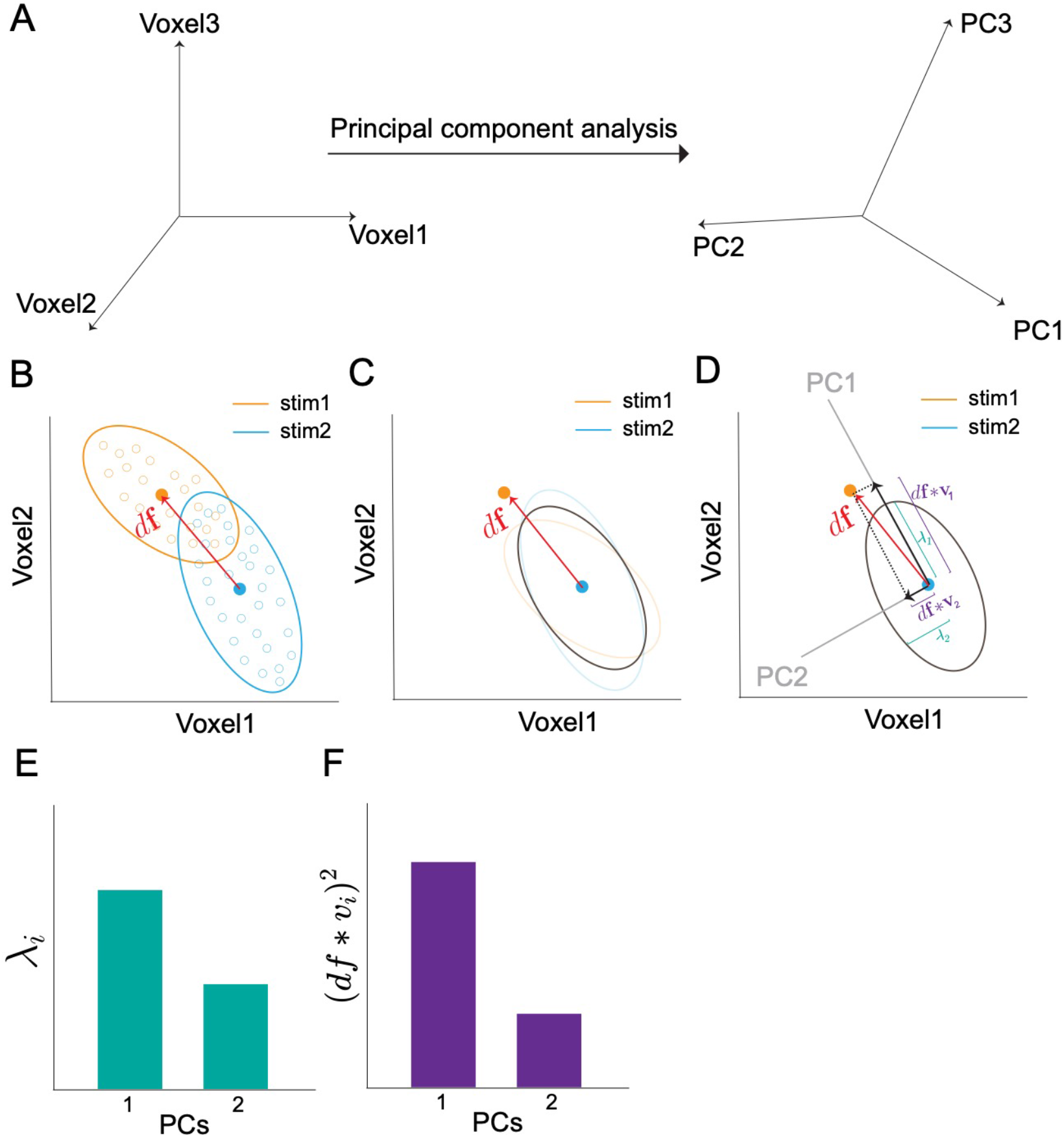
Schematics of eigen-decomposition of information. ***Panel A*** depicts that the information contained by trial-by-trial high-dimensional population responses can be calculated in the eigenspace (obtained by principal component analysis) instead of the original voxel space (i.e., Eq. 12). Thanks to the linear independence of the eigenspace, the information of the whole population can be simply reformulated as the summation of information along each eigendimension (i.e., 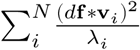, see Eq. 12 in Methods and Materials). ***Panel B*** depicts two 2d response distributions of two voxels towards two stimuli. The yellow and blue ellipses also show the direction of covariances (i.e., **Q**_**1**_and **Q**_**2**_). The red vector (*d***f)** can be viewed as the Euclidean distance between the two distributions (see. Eq. 2 with the assumption of *ds*=1). The gray ellipse in *Panel C* depicts the averaged covariance **Q** (i.e., averaged **Q**_**1**_ and **Q**_**2**_, see Eq. 3). ***Panel D*** depicts that the averaged covariance can be decomposed into two principal components (PC1 and PC2). The variance λ_*i*_ along each PC is illustrated in ***panel E***. Intervoxel RCs result in a larger variance in PC1 (i.e., λ_1_) than PC2 (i.e., λ_2_). The squared projected signals (*d***f** * **v**_*i*_)2 on each PC are illustrated in ***panel F***. Note that the sum of squared projected signals (i.e., the sum of bars in ***panel F***, 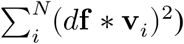 is a constant, which amounts to the norm of the signal vector *d***f**.

The intuition of this formulation can be illustrated in the two-voxel scenario in Fig. 7. In Fig. 7A, the two voxels have substantial RCs and the majority of the variance lies in the 1^st^ eigendimension (i.e., principal component, PC). Here, λ_*i*_ indicates the amount of variance along with the *i*-th PC, and *d***f** * **v**_*i*_ indicates the squared projection of the signal vector (*d***f**) onto that PC. The information on that PC is equal to the ratio between the squared projected signals and variance 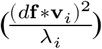. Obviously, the larger the projected signals *d***f** * **v**, the smaller variance λ, the more information will be on a PC. Furthermore, the total information in a population is the sum of information across all eigendimensions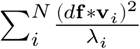. This formulation allows us to disentangle the contributions of projected signals and variance on each eigendimension. This method of information decomposition has been recently proposed and proven to be sensitive when a relatively small number of units and trials are analyzed 25,32.

We first performed the analysis of information decomposition to examine the effects of tuning-compatible RCs in the neuron-encoding model. Specifically, we calculated the eigenvalue (λ_*i*_, Fig. 8A), the squared projected signal ((*d***f** * **v**_*i*_)2, Fig. 8B), their ratio (i.e., the information on this PC, 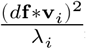, Fig. 8C) on each PC, the cumulated information across the first few PCs (Fig. 8D), and the total information (Fig. 8E), as a function of RC strength. We found that the variance of population responses is concentrated on the first few PCs if RC strength increases (Fig. 8A). This is not surprising because increasing RCs must produce a low-dimensional manifold along which population activity fluctuates. Interestingly, increasing tuning-compatible RCs also heightens the projected signals ((*d***f** * **v**_*i*_)2) on the first few PCs. More projected signals on high-variance PCs result in fewer signals on low-variance PCs. Given that both the projected signal ((*d***f** * **v**_*i*_)2), as the numerator, and the variance λ_*i*_, as the denominator, decrease significantly, it is difficult to predict the changing direction of their ratio 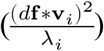 by intuition. By plotting 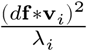, we found that tuning-compatible RCs reduce information on high-variance PCs but enhance information on low-variance PCs (Fig. 8C). The former effect is much more dominant, resulting in less total information in the population (Fig. 8D). In other words, the detrimental effects of tuning-compatible RCs in neurons (as shown in Fig. 8E) are primarily driven by information reduction on high-variance PCs.

**Figure 8.**
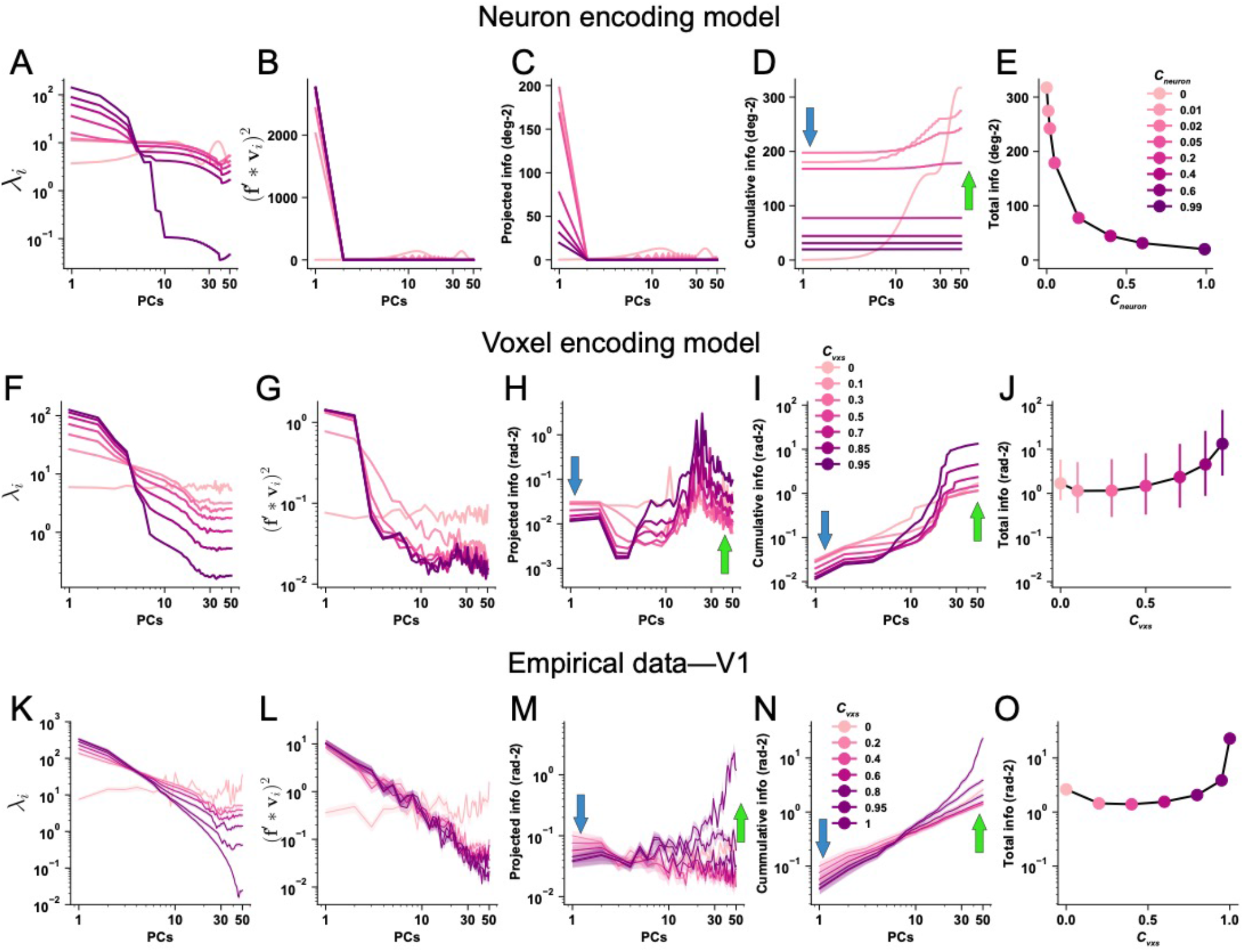
Eigen-decomposition information analyses for the effects of tuning compatible response correlations in the neuron-encoding model (***Panels A-E***), the voxel encoding model (***Panels F-J***), and empirical data of V1 (***Panels K-O***). In ***Panels A-D, Panels F-I***, and ***Panels K-O***, the x-axes are PCs ranked by their variance from high to low. ***A***. eigenvalue (i.e., variance λ_*i*_) on each PC. The variance of population responses is heightened on the first few PCs and reduced on the last few PCs when RC strength increases. ***B***. squared projected signals (*d***f** * **v**_*i*_)2 on each PC. As RC strength increases, the projected signals also increase on the first few PCs. More projected signals on the first few PCs will inevitably lead to fewer projected signals on other PCs. ***C***. information 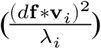 on each PC. ***D***. cumulative information of the first few PCs (i.e., the sum of the first few data points in ***Panel C***). Information quickly saturates given the presence of tuning-compatible RC. Without RC, information slowly increases but eventually reaches a higher level. The blue and green arrows highlight the antagonistic information change on high- and low-variance PCs, respectively. ***E***. information as a decreasing function of RC strength. Note that the data points in ***Panel E*** correspond to the last data points (i.e., PC = 50) of the curves in ***Panel D. Panels F-J*** depict the results of the same analyses for tuning-compatible RCs in the voxel encoding model. The error bars in ***Panel J*** represent the 95% confidence intervals of 100 independent simulations. The key difference here is the information enhancements on low-variance PCs are much more pronounced in voxels as compared to neurons, producing a U-shaped function (***Panel J***). These effects are further validated in the empirical fMRI responses in human V1. For each line in ***Panels K-O***, we calculated 72 independent samples (6 subjects x 2 hemispheres x 6 pairwise comparisons). There are 6 pairwise comparisons because of the four stimulus orientations. The solid lines and the shaded areas represent the mean and the S.E.M of the samples. The error bars are very small and can be barely seen in ***Panel O***.The conventions of the arrows are kept in subsequent figures.

We then applied the same analyses to examine the effects of tuning-compatible RCs in the voxel-encoding model. Similar to neurons, tuning-compatible RCs among voxels induce higher variance (Fig. 8F) and higher projected signals (Fig. 8G) on the first few PCs. It is thus nontrivial to predict the changing trend of their ratio 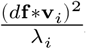. Again, we found that tuning-compatible RCs also reduce and enhance information on high- (see the blue arrow in Fig. 7H) and low-variance PCs (see the green arrow in Fig. 8H), respectively. Different from the scenario of neurons, these two antagonistic effects together produce a U-shaped function (Fig. 7I&J). Also, the information enhancements on low-variance PCs dominate, eventually resulting in an overall larger amount of information (Fig. 8I&J).

We further performed the same analyses on the empirical fMRI data in V1. We found similar results as predicted by the voxel-encoding model. Increasing RCs inevitably produces a few high-variance PCs (Fig. 8K) and stronger projected signals on these PCs (Fig. 8L). Although RCs lessen the information on those high-variance PCs (see blue arrows in Fig. 8M&N), they also disproportionally enhance information on low-variance PCs (green arrows in Fig. 8M&N), producing a U-shaped function of total information (Fig. 8O). These results are consistent with the voxel-encoding model above. Analyses in V2 and V3 show highly similar results to those in V1 (see Supplemental Information Fig. S1). As a control, we also performed the information decomposition analysis on the neuron- and voxel-encoding models assuming tuning-irrelevant RCs. Results showed that tuning-irrelevant RCs substantially attenuate the information reduction effect on high-variance PCs, and overall enhanced information mainly comes from the information enhancements on low-variance PCs (see Supplemental Information Fig. S2).

In sum, despite the seemingly drastically opposite effects of tuning-compatible RCs in neuronal and voxel populations, we found a unified mechanism underlying the two scenarios. In both types of data, increasing tuning-compatible RCs produces two antagonistic consequences: information reduction on high variance PCs and information enhancements on low-variance PCs. The relative balance of these two effects determines the exact shape of the information function as RC strength increases, because a continuum of possible effects (i.e., monotonic increasing/decreasing, U-shaped function) may occur. For neurons, the effect of information reduction dominates, and therefore tuning-compatible RCs monotonically reduce information. For voxels, the effect of information enhancement becomes more pronounced and therefore produces the U-shaped function of information. These effects are further validated in the empirical fMRI data.

These results also render the interpretations of improved MVPA accuracy more complicated. We know linear Fisher information is monotonically related to MVPA accuracy and sensory information is a U-shape function of RC here. Enhanced sensory information (i.e., improved MVPA accuracy) thus can arise from many possibilities. For example, both the increase and reduction of RC can lead to enhanced information as long as the baseline condition lies at the ridge part of the U-shaped function. It remains largely unclear how cognitive factors modulate population response properties in fMRI. More broadly, the brain can attain a better coding scheme by modulating tuning, noise, and response correlations, or combinations of all these factors, a much more flexible mechanism than previously believed. This implication also invites deeper consideration of previous empirical studies using MVPA accuracy to characterize brain functions.

### U-shaped information function in larger voxel pools

It has been shown that detrimental response correlations may masquerade as beneficial given noisy measurements and a limited number of neurons in an empirical study (see Fig. 9A) 2. We further systematically modeled the effects of correlation strength and pool sizes. As shown in Fig. 9C, the U-shaped function held when the number of voxels increased up to 2000, which is much larger than the number of voxels in a typical MVPA analysis. These results are also consistent with our past theoretical work 26. We cannot completely exclude the possibility that sensory information will eventually saturate at a lower asymptote level when pool size goes infinite (i.e., the signature of information-limiting correlation). But our results should hold for the majority of MVPA fMRI studies.

**Figure 9.**
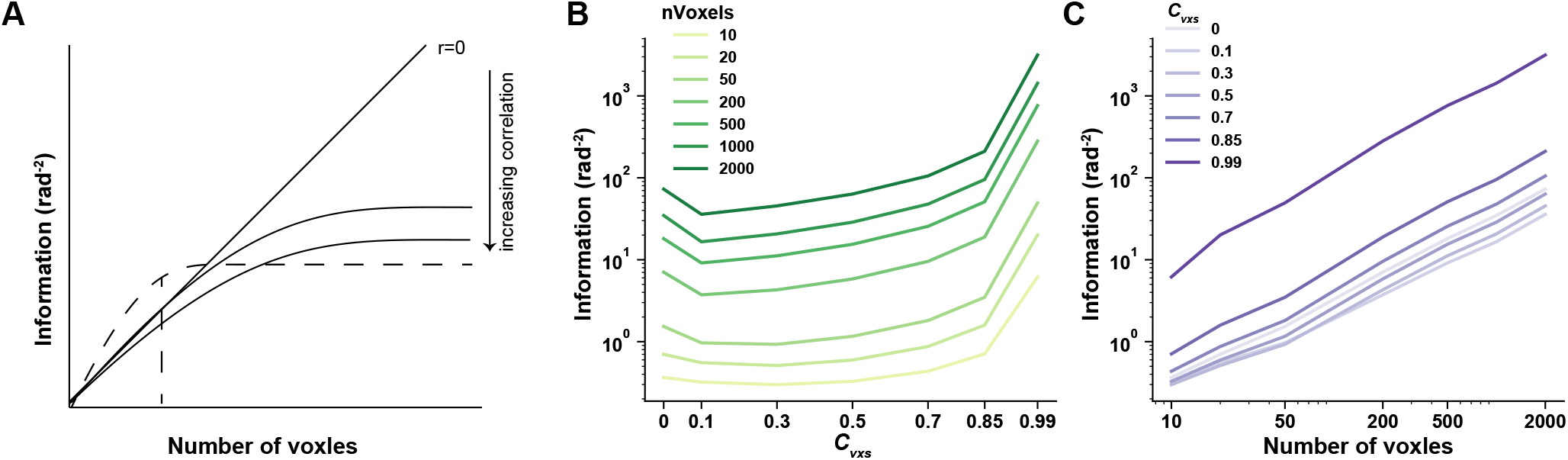
the effect of pool size on stimulus information. Theoretically detrimental neural correlations may manifest as beneficial due to noisy measurements in limited pool size. In ***Panel A***, the solid lines are the stimulus information as a function of pool sizes (i.e., the number of voxels). Increasing correlations dampens the stimulus information and stimulus information will eventually saturate given the presence of correlation. The dashed line indicates stimulus information calculated in empirical data. Correlations may masquerade as beneficial (i.e., the dashed line is higher than the diagonal line for small pool size). We further calculated stimulus information as a function of correlation strength (***Panel B***) and pool sizes (i.e., number of voxels, ***Panel C***). Results showed that the U-shaped function is ubiquitous for increasing pool sizes and information does not saturate up to 2000 voxels, which is far more than the number of voxels used in MVPA analyses in a typical empirical study.

## DISCUSSION

The majority of empirical studies in neurophysiology have demonstrated the detrimental role of neuronal RCs. Weakened interneuron RCs have been found to accompany improved behavioral performance in many cognitive tasks. Here, we investigate the effects of RCs in fMRI using an information-theoretic approach. We compare the bias-corrected linear Fisher information between the data where RCs are intact or hypothetically removed. We find higher stimulus information in the former case, supporting the beneficial role of RCs in fMRI data. We then systematically manipulate RC strength and find that stimulus information follows a U-shaped function of RC strength. These results stand in stark contrast to the monotonically detrimental effects of RCs documented in neurophysiology. Interestingly, this discrepancy between neurophysiology and fMRI can be well explained by a voxel-encoding model that bridges neuronal and voxel activity. Most importantly, information decomposition analyses further suggest that RCs reduce information on high-variance PCs, and in the meantime, enhance information on low-variance PCs. The two antagonistic effects together may result in increasing, decreasing, or U-shaped information functions, explaining a wide range of theoretical and empirical findings. This information decomposition approach also highlights the complexity of quantifying information and can serve as a unified mathematical framework that helps resolve debates in computational neuroscience.

### The effects of response correlations in neurophysiology

The effects of RCs on stimulus information have been a matter of debate over the decades 2. One major contribution of our work is to use a unified framework—information decomposition—to quantify stimulus information in fMRI data. This method in theory can be used on different modalities (e.g., neuronal spikes, fMRI, EEG sensors). Our analyses demonstrate two non-trivial issues when quantifying information in a population. First, Eq. 12 reveals that the information on an eigendimension depends on the projected signals (*d***f** * **v**_*i*_) and variance (λ_*i*_) on that dimension. As RC strength increases, both the squared projected signals (*d***f** * **v**_*i*_)2, as the numerator, and the variance (λ_*i*_), as the denominator, exhibit decreasing functions across different PCs. The information on each PC, as their ratio (i.e 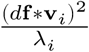), theoretically cannot be easily predicted by intuition and should be calculated formally. Second, even though the information changes on individual eigen-dimensions are carefully quantified, it is still difficult to infer the total information because RCs lessen information on high-variance PCs but enhance information on low-variance PCs. These two antagonistic effects together in theory could result in any shape of information function, depending on their relative strength.

It has been shown that only a specific type of correlation—”differential correlations” limit information but their presence might be hard to detect due to the limited number of trials and neurons recorded in empirical data 33. The information decomposition analysis has been recently proposed as a feasible solution to circumvent this limitation and help detect “differential correlations” using a much smaller number of units and trials 25. Instead of the reduced information on high-variance PCs, we find a significant contribution of enhanced information on low-variance PCs, which is usually ignored in previous literature. This phenomenon indicates that careful quantification of covariance structure in fMRI data and formal calculation of information are needed in future empirical studies.

Our results speak to several hassles in previous theoretical work. Previous studies have investigated several factors that may mediate the effects of RCs. For example, a bulk of theoretical studies have investigated how tuning heterogeneity, the magnitude and form of RCs, and behavioral relevance modulate the effects of RCs 1,2,5-7. We want to emphasize that there exist very complex interactions between these factors and the exact values of parameters used in theoretical modeling determine the result. For example, it has been shown that tuning-compatible RCs among neurons may turn out to be beneficial in a population with very heterogeneous tuning curves 34. The complexity of analyzing information necessitates investigations on a systematic and wide range of parameters to help comprehend the interactions between these factors. Moreover, future empirical studies need to thoroughly measure these parameters, and researchers need to perform formal information-theoretic analyses in future work.

### The comparisons between MVPA and information-theoretic approaches

Fisher information is tightly linked to MVPA decoding accuracy because the inverse of Fisher information is defined as the lower bound of (co)variance of an unbiased maximum likelihood decoder 35. In this paper, we used the information-theoretical approach instead of the conventional decoding approach because it is advantageous in terms of investigating the effects of RC.

Both approaches have their respective advantages and disadvantages. Decoding (e.g., support vector machine) is powerful when data contain a large number of dimensions (i.e., voxels) but only a few samples (i.e., trials), a scenario that most fMRI studies encounter. Also, decoding usually employs a discriminative modeling approach requiring no assumption of the generative distributions of fMRI data. Moreover, decoding is suitable for almost any feature dimension regardless of whether the stimulus variable is continuous (e.g., orientation, direction) or discrete variables (e.g., object categories). However, decoding only provides a single accuracy value without showing sufficient details of representational geometry 36. In particular, decoding falls short to disentangle the effects of signal, noise, and response correlations on population codes (e.g., differentiate the cases in Fig. 2C&D).

By contrast, linear Fisher information has been widely used in computational neuroscience to demonstrate the quality of population codes in neural spike data. Linear Fisher information is based on the generative mechanism of data because by definition it is related to maximum likelihood decoding and thus guaranteed to be statistically optimal. However, using a limited number of units and trials may yield biases in the estimation of linear Fisher information. The analytical solution to correct the bias is based upon the assumption of Gaussian variability, and violations of Gaussian variability in empirical data might introduce additional bias.

In this study, we only consider one visual feature (i.e., orientation) and human early visual areas where MVPA has been extensively employed in literature 20,21. But our approach here is widely applicable and future studies might need to test the effects of response correlations in more visual features (e.g., motion direction) and sensory domain. In sum, we contend that both MVPA and the information-theoretic approach are fruitful in future fMRI research.

### The interpretation of linear Fisher information

Fisher information has an explicit physical meaning. The information can be converted to the discrimination threshold of an optimal linear decoder. As such, linear Fisher information is often utilized as a reference to infer the readout efficiency from neural responses to behavior. If a behavioral threshold is close to that indicated by Fisher information, it suggests that downstream decision units efficiently extract nearly all information in neural responses to guide action. This possibility has been recently confirmed in ref. 25. Otherwise, if the behavioral threshold is higher than that indicated by Fisher information, it suggests a significant information loss from neural responses to behavior.

We want to specially emphasize that this information-behavior comparison in typical unit-recording studies is inappropriate in fMRI due to the intrinsic low signal-to-noise ratio of fMRI data. Unlike neurophysiology, decoding accuracy in fMRI rarely reaches human behavioral performance on the same stimuli. For example, to discriminate two high-contrast gratings with orthogonal orientations, decoding accuracy on fMRI data can only achieve ∼70%-80% accuracy but human behavior can easily reach nearly 100% 20,21. Here, we obtained ∼1 rad^-2^ information

(as shown in Fig. 3), and this corresponds to the ∼77º orientation discrimination threshold given the 75% accuracy. This is consistent with classical fMRI decoding results 20,21 but far worse than the human behavioral threshold of orientation discrimination (∼1-2º). Like MVPA, we can only compare the relative change of linear Fisher information across different cognitive states (e.g., attention vs. unattention).

The low signal-to-noise of fMRI data also endows a potential limitation of this study. Here, large orientation differences (i.e., 45 and 90) are used in the fMRI experiment and for the calculation of linear Fisher information. This is indeed consistent with the majority of decoding studies in fMRI. However, Fisher information by definition only measures the local curvature of log-probability distributions, and neurophysiological studies typically use a pair of stimuli with small differences (e.g., < 10º). Theoretically, the calculation of the covariance matrix via Eq. 3 may be imprecise when a pair of stimuli with large disparities are used. We argue that this should not be a problem for fMRI because (1) the signal-to-noise ratio is intrinsically low in fMRI signals, and (2) Fig.3b shows that the empirically measured covariance matrices for different stimuli with large disparities are highly similar (i.e., stimulus-invariant covariance matrices). Eq. 3 is therefore appropriate here. The calculation of linear Fisher information in the neuron-encoding model may be imprecise (Fig. 5C) but this does not affect the conclusion as here we only focus on the qualitatively detrimental effects of tuning-compatible noise correlations (i.e., the decreasing line in Fig. 5C) in neuronal populations, a phenomenon that is consistent with the case of stimuli with slight differences.

Like neurons, voxel activity varies substantially across trial-by-trial. Such trial-by-trial variability reflects underlying neuronal variability as well as many other sources of noise. There exist at least three types of noise that can cause the variability of voxel activity. First is the measurement noise related to fMRI data acquisition, such as thermal noise, electronic noise, etc. These types of noise are unlikely related to voxel tuning and thus unlikely produce tuning-compatible RCs. One exception is head motion because head motion might induce structured correlations between nearby voxels and this effect may act as tuning-compatible RCs. It remains an active field of research in fMRI how to best control and minimize the influences of head motion during data acquisition and processing 37. The second type of noise is related to some global brain functions (e.g., arousal) and this type of noise is unlikely related to voxel tuning either. The third type of variability comes from underlying neuronal variability in a voxel and should produce tuning-compatible RCs as we show analytically and empirically above. This type of variability is most intriguing to neuroscientists because it usually reflects important aspects of stimulus 2,19. Unfortunately, current fMRI technology does not allow us to distinguish different sources of noise, and it remains a challenge for future fMRI studies to dissect the effects of noise on empirical data.

Here, we found that only by incorporating tuning-compatible RCs in the voxel-encoding model we can reproduce the U-shaped function observed in the real data. The two assumed forms of RCs (i.e., tuning-compatible and tuning-irrelevant RCs) here are certainly oversimplified descriptions of real RCs. In empirical data, RCs are likely to be a mixture of both types. The small differences between the results from simulated voxels (Fig. 7F-J) and real voxels (Fig. 7K-O) might due to the mixed nature of RCs in empirical data.

Moreover, the definition of RC also varies across subfields of fMRI. In this paper, we follow the standard definition of RC in neurophysiology and define it as the trial-by-trial activity correlation between two voxels, or “Beta-series correlation” in fMRI literature 15. But some studies also regard resting-state functional connectivity or task-based functional connectivity 38 as response correlation. These three types of correlations are by definition different but might be quantitatively related in empirical data. Future studies are needed to further test their links.

### Implications for future fMRI studies

What are the implications for future fMRI practice? We would like to emphasize three possible aspects where our framework is useful.

First, in this study, we manipulated the strength of RC when calculating linear Fisher information of stimuli. This is only a data analysis method and by no means suggests that one can easily manipulate RC strength in empirical data. In a standard decoding study, there is no need to formally quantify RCs or their effects if one only focuses on decoding accuracy across different cognitive states or across brain regions (e.g., searchlight analysis). One practical suggestion is that one should use the decoding method that takes into consideration the correlation structure in data if RCs in general improve decoding. For example, it has been shown that support vector machine or logistic regression that acknowledges voxelwise RCs outperform a nearest-neighbor decoder that does not consider RCs 39. Also, some recent efforts on Bayesian decoding require explicit modeling of the correlation structure. Thus, measuring and quantifying voxel correlations is an essential step toward more robust brain decoding 27.

Second, even though many factors (e.g., acquisition noise) may result in voxel RCs and these RCs may help decoding, we still want to carefully control these factors during fMRI data acquisition and processing. This is because 1) they may lower the signal-to-noise ratio of individual voxels, and 2) more importantly, lead to inaccurate neuroscientific interpretations of fMRI data 40.

Third, researchers cannot directly manipulate RCs in empirical data but can manipulate cognitive states of the brain, and different cognitive states might in turn alter RC structures. Despite the profound evidence of top-down modulations on RCs in neurophysiology 13, it remains unclear whether altered RC structures underpin cognitive processes in humans. For example, attention and perceptual training have been shown to enhance decoding accuracy in human visual cortex 23,41. But it remains unclear whether the enhanced decoding accuracy arises due to altered RCs. The considerable effects of RCs on stimulus information at least suggest that modulating RCs is an effective way to alter stimulus information in multivariate fMRI data. In particular, in this paper we isolate the effects of RCs by keeping other aspects of voxel responses intact. In realistic experiments, cognitive processes (e.g., attention) can alter signals, noises, RCs, or combinations of these factors. It is the interactions between these factors that produce the outcome stimulus information. Future studies are needed to further dissect the computational mechanisms of altered decoding accuracy in human fMRI.

## MATERIALS AND METHODS

### fMRI experiment

#### Stimuli and task

We analyzed the fMRI data collected by ref. 42. The datasets are publicly available on the openNeuro website (https://openneuro.org/datasets/ds000113/versions/1.3.0). Six subjects participated in the study. Briefly, two flickering sine-wave gratings (0.8-7.6º eccentricity, 160º angular width on each visual hemifield with a 20º gap on the vertical meridian) were presented on both sides of the fixation point. The orientations of the two gratings varied independently across trials. The orientations were drawn from 0º, 45º, 90º, and 135º with equal probabilities. A stream of letters was presented at the center-of-gaze throughout each scanning run. Subjects were instructed to main their fixation and perform a reading task. To ensure subjects’ task engagement, subjects were tested on a question related to the reading text at the end of each run.

On each trial, an orientation stimulus lasted 3 s and was followed by a 5 s blank. Each scanning run consisted of 30 trials. The 30 trials also included 10 randomized blank trials. The 1st trial could not be a blank trial, and there were no two consecutive blank trials. Blank trials could appear on either side while the grating on the other side was intact. Each subject completed 10 scanning runs.

#### MRI data acquisition and processing

A T2*-weighed echo-planar imaging (TR/TE=2000/22ms) sequence was used to acquire fMRI data. In the original experiment, the subjects were scanned at four different resolutions. Here we only analyzed the 2-mm isotropic data because they gave the best decoding accuracy 43. 121 functional volumes were acquired in each run (FoV=200mm, matrix size 100 × 100, 37 slices, GRAPPA accel. factor 3). The echo-planar images covered the occipital and parietal lobes. Each subject was also scanned for a high-resolution T1-weighted image (0.67mm isotropic). Additionally, we conducted standard retinotopic scans to define low-level visual areas.

The pial and white surfaces were reconstructed based on the high-resolution T1-weighted images using the standard FreeSurfer pipeline. All functional volumes underwent slice-timing correction, motion correction, and registration to T1 images. Retinotopic data were analyzed using the 3dRetinophase tool in AFNI to generate polar angle and eccentricity maps on cortical surfaces. Bilateral V1-V3 were defined on spherical cortical surfaces based on polar angle and eccentricity maps. The vertices in each ROI on cortical surfaces were then transformed back to EPI space using the AFNI 3dSurf2Vol function to locate corresponding voxels.

We used AFNI 3dDeconvolve function to build general linear models (GLM). Particularly, the “-stim_times_IM” option of the function was used to model each presentation of stimuli (i.e., trial) as an independent predictor. We implemented two GLMs separately for two hemispheres due to independent stimuli presentations for each hemisphere, and each GLM only included the stimuli presented contralaterally. In each GLM, demeaned head motion parameters, and constant, linear, and quadratic polynomial terms were also included as nuisance predictors. In each subject, we concatenated the time series of 10 scanning runs and fitted a single GLM to the combined data.

### Calculation of linear Fisher information in population responses

In computational neuroscience, Fisher information is a standard metric that assesses the amount of information carried by observed neuronal responses **r** with respect to stimulus variable *s*. Two signatures of **r** should be noted. First, the neuronal response **r** is high-dimensional (i.e., a vector) because it represents the responses of many neurons or voxels. Second, due to trial-by-trial variability, the **r** across trials follows a multivariate response distribution in the high-dimensional space. If the response distribution belongs to the exponential family with linear sufficient statistics (i.e., information can be optimally decoded via a linear decoder), Fisher information can be further simplified as linear Fisher information 44. Henceforth, we call it *information* for short.

Suppose that, in an experiment, we attempt to discriminate two stimuli *s*_*1*_ and *s*_2_, based on the measured responses of *N* voxels in *T* trials for each stimulus. This is a typical binary classification scenario in fMRI 20,21. Here Fisher information can be understood as to what extent the two stimuli can be discriminated based on population responses.

In the ideal case that we have an infinite number of trials *T*, the information delivered by population responses can be written as:

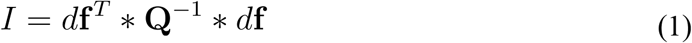

where *d***f** and **Q** are defined as:

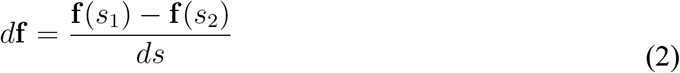

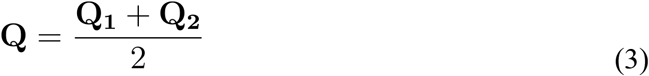

**f** (*s*_1_) and **f** (*s*_2_) are vectors, representing the mean responses across trials of the neuronal population for stimuli *s*_1_ and *s*_2_, respectively. *ds* is the absolute stimulus difference (i.e., *ds* = |*s*_1_ − *s*_2_|). **Q**_**1**_ and **Q**_**2**_ are the covariance matrices of the population responses towards stimuli *s*_1_ and *s*_2_, respectively. Note that Eq. 1 has no assumption about Gaussian response variability.

Eq.1 has several important implications. First, stimulus information depends on 1) the Euclidean distance between the two high-dimensional response distributions (i.e., **f** (*s*_1_) − **f** (*s*_2_)), (2) the covariance (i.e., the variance of individual voxel response and voxelwise response correlations) of the population (i.e., **Q**_**1**_ and **Q**_**2**_), and (3) the interaction between (1) and (2). We illustrate this issue in Fig. 2. Eq.1 also highlights the complexity of quantifying information as any effort that merely considers one factor might fail to yield meaningful results because all factors (i.e., mean responses, variance, and response correlations) and their interactions matter.

Any real experiment, however, must have a finite number of trials *T*. The estimations of the statistical properties (e.g., **Q**_**1**_ and **Q**_**2**_) of population responses therefore might be imprecise. Imperfect estimations of these statistics can lead to potential biases in the estimation of information. This is especially common in the field of fMRI because the number of trials sampled in an experiment is usually smaller than the number of voxels. ^Kanitscheider, et al. 24^ have shown that the bias induced by imperfect estimations of response properties indeed exists but fortunately can be corrected analytically. The bias-corrected linear Fisher information (*I*_*bc*_) can be written as:

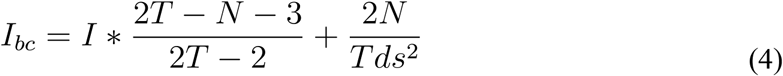

where it has been also proved that the bias manifests differently in populations with and without response correlations 24. In the latter case, voxel responses are independent, and this is equivalent to shuffling the responses of each voxel separately across many trials, a popular strategy used in literature to investigate the effect of neuronal response correlations. In this case, linear Fisher information (*I*_*shuffle*_) can be corrected by:

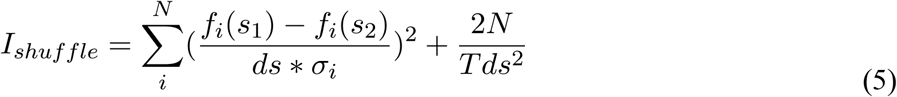

where *f*_*i*_(*s*_1_) and *f*_*i*_(*s*_2_) are the mean responses of the *i*-th voxel across trials. 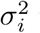 is the averaged variance (i.e., 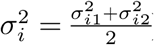), where 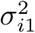and 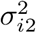 are the variance of the responses of *i*-the voxel across trials. We used Eq. 4 to calculate stimulus information in empirical data (puce bars in Fig. 3B), and Eq. 5 to calculate stimulus information when RCs are removed (gray bars in Fig. 3B). Note that although linear Fisher information *per se* (Eq. 1) has no assumption of Gaussian variability, the analytical solution for bias correction assumes Gaussian variability (Eqs. 4&5). But it has also been shown that this bias correction solution is also robust to non-Gaussian cases 24. Also, the bias-correction method is only valid when *T > (N+2) / 2*. We thus chose N = 50 in all neuron- and voxel-encoding modeling below and also when analyzing empirical fMRI data.

For each orientation, we defined the information of each orientation as the averaged information of pairwise discriminating that orientation and the other three orientations. For example, the information of 0° is defined as 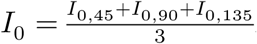.

### Multivariate Gaussian variability of trial-by-trial voxel responses

Linear Fisher information has no assumption about Gaussian variability of trial-by-trial responses. The analytical solution of bias correction given a limited number of units and samples, however, rests upon the assumption that the trial-by-trial population responses follow a multivariate Gaussian distribution. Although Gaussian variability of voxel responses has been either implicitly assumed or explicitly formulated in several previous studies 27,31, it remains a presumption rather than a grounded finding.

We tested whether trial-by-trial population responses follow multivariate Gaussian distribution in our data. For each stimulus, the data can be represented as an *N* (voxels) x *T* (trials) matrix (Fig. 1B). Here voxels and trials can be viewed as features and samples, respectively. We performed the Henze-Zirkler multivariate normality test (implemented by the *multivariate_normality* function from the *pingouin* python package) on the data matrix. The Henze-Zirkler test has good power against alternatives to normality and is feasible for any dimension and sample size 45. We performed the Henze-Zirkler multivariate normality test for the voxels in each ROI, hemisphere, and subject. We found no significant violation (all p-values > 0.05) of multivariate Gaussian variability in a total of 144 tests (6 subjects x 2 hemispheres x 3 ROIs x 4 stimuli), revealing multivariate Gaussian distributions as an appropriate approximation of trial-by-trial voxel responses.

### The titration approach to derive information as a function of response correlation strength

We parametrically manipulated the strength of RCs between voxels. We multiplied the off-diagonal items in the covariance matrices **Q**_**1**_ and **Q**_**2**_ with a coefficient *c*_*νxs*_ but kept the diagonal items intact. For the item (*q*_*ij*_) at the *i*-th row and the *j*-th column in **Q**_**1**_ or **Q**_**2**_, we have:

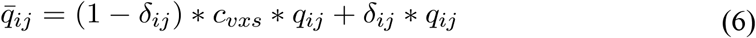

where *δ*_*ij*_ is the Kronecker delta (*δ*_*ij*_=1 if *i*=*j* and *δ*_*ij*_= 0 otherwise). We then used the resultant matrices 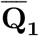 and 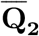 instead of **Q** or **Q** to calculate information using Eqs. 1-3. By this method can we titrate the strength of RC and investigate how it impacts information. Note that the two scenarios (i.e., with and without RC) in Fig.1 correspond to the condition that *c*_*νxs*_ = 1 and 0, respectively. Here we did not use Eqs. 4&5 to calculate bias-corrected linear Fisher information because (1) it is unclear the bias when *c*_*νxs*_ is within [0, 1], and (2) we are interested in the overall changing trend of information as a function of RC strength rather than the absolute magnitude of information.

### Neuron- and voxel-encoding models

#### Neuron-encoding model

We simulated 50 orientation-selective neurons whose preferred orientations equally span within (0, π]. The von Mises tuning curve of the *k*-th neuron is:

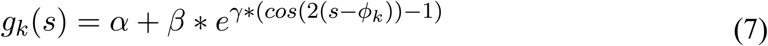

where *α, β, γ* control the baseline, amplitude, and tuning width and we set them as *α* = 1, *β* = 19, *γ* = 2 34. *φ*_*k*_ is the preferred orientation of that neuron. We assume that neuronal spikes follow Poisson statistics (i.e., variance equal to mean firing rate).

Two types of RC structures were investigated here. The first one is *tuning-compatible RC* (TCRC), indicating that the RC between a pair of neurons is proportional to the similarity of their tuning functions:

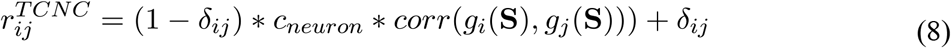

where 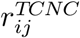 is the tuning-compatible RC between the *i*-th and the *j*-th neurons. *g*_i_ (**S**) and *g* (**S**) are tuning functions of the two neurons. *δ*_*ij*_ is the Kronecker delta (*δ*_*ij*_ =1 if *i=j* and *δ*_*ij*_ =0 otherwise). **S** is the vector of all possible orientations within (0, π]. We denote **R**^TCRC^ as the tuning-compatible RC matrix. The theoretical derivation and empirical measurement of tuning-compatible RC will be introduced in later sections.

We also investigated a control type of RCs called *tuning-irrelevant response correlation* (TIRC). We shuffled the pairwise RCs in **R**^TCRC^ such that the rows and columns were rearranged in a random order but the diagonal items kept unchanged. We denote the resultant matrix as **R**^TIRC^. By this shuffling method, the overall distribution of the items in **R**^TCRC^ and **R**^TIRC^ are identical but the pairwise RCs in **R**^TIRC^ no longer resemble the tuning similarity between two neurons.

We also applied the ‘titration’ approach (response correlation coefficient *c*_*neuron*_) to manipulate the strength of RCs in both cases and calculated the linear Fisher information of discriminating orientations 0 and 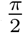 in the neuronal population. For the case of tuning-compatible RC, the information is deterministic because **R**^TCRC^ is fixed; for the case of tuning-irrelevant RC, we created **R**^TIRC^ and calculated information 100 times.

#### Voxel-encoding model

The voxel-encoding model is built based on the neuron-encoding model. Given the 50 orientation-selective neurons, we further simulated 50 voxels using the voxel-encoding model:

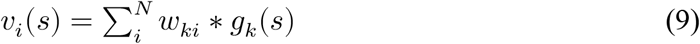

where *ν*_*i*_(*s*) is the tuning curve of the *i*-th voxel, and *w*_*ki*_ is the connection weight between the *k*-th neuron and the *i*-th voxel. The weighting matrix **W** maps neuronal tuning curves to voxel tuning curves. Instead of Poisson noise, we assumed additive noise on voxel activity the response variance 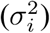 of each voxel is drawn from a Gamma distribution (i.e., 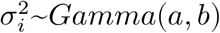), where *a* = 1.5 and *b* = 4 are the scale and the shape parameters corresponding to the Gamma distribution with mean = 6, and variance = 24.

Similar to the neuron-encoding model, we also assumed tuning-compatible response correlations between voxels:

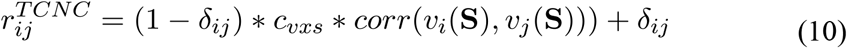

As a control, we also assumed tuning-irrelevant response correlations 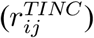 by shuffling rows and columns of **R**^TCRC^ as described above.

We manipulated *c*_*νxs*_ and calculated the information under the regimes of tuning-compatible RCs and tuning-irrelevant RCs (Fig. 5D). Because the voxel tuning curves and response variance depend on the weighting matrix **W** (Eq. 9) and the Gamma distribution, respectively. We run the simulation 100 times, and in each simulation, a new weighting matrix **W** is created by:

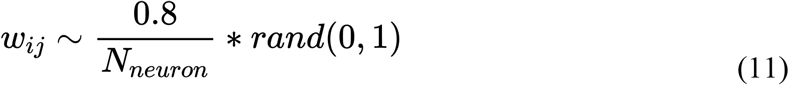

where *N*_*neuron*_ is the number of hypothetical neurons (i.e., 50 in our case) in the voxel-encoding model. We set the scaling factor 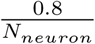 to ensure the voxel activity was mostly below 5%, which is consistent with the percent bold signal change in real fMRI data.

### Eigen-decomposition of information

Eq. 1 can be further reformulated:

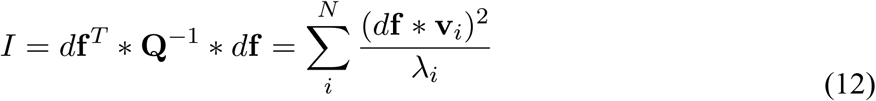

where **v**_*i*_ and λ_*i*_ are the *i*-th unit eigenvector and its corresponding eigenvalue. *d***f**, the signal vector, is defined in Eq. 2. The *d***f** * **v**_*i*_ can be viewed as the projected signals on the *i*-th eigendimension. λ_*i*_, the eigenvector, indicates the variance on the *i*-th eigendimension. Thus 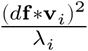 can be considered as the signal-to-noise ratio, or information, on the *i*-th eigendimension. Because eigendimensions are linearly uncorrelated, the total information should be the sum of information across all eigendimensions.

### Theoretical derivation of tuning-compatible response correlations in voxel populations

To derivate the theoretical feasibility of tuning-compatible RCs in voxel populations, we simulated the RC matrix using the voxel-encoding model 1000 times. In each simulation, we randomly generated the weighting matrix **W** (using Eq. 11) and only considered the variability propagated from the neuronal to the voxel level. Tuning curves of voxels can be computed based on **W** (Eq. 9), and the tuning similarity between the *i*-th and the *j*-th voxels can be computed by 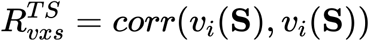. Note that here we did not assume any RCs at the neuron level as that in the neuron-encoding model above. For a given stimulus *s*, due to Poisson variance, the covariance matrix (**Q**_*neuron*_) of neuronal responses in Eq. 1 should be: **Q**_*neuron*_ = *diag*(***g***(*s*)). We can analytically derive the covariance (**Q**_*νxs*_) of voxel responses as:

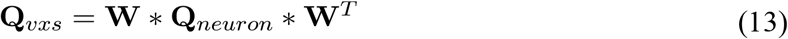

And the RC matrix 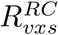 can be calculated based on the covariance **Q**_vxs_. Given 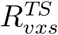 and 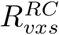, we calculated the correlation between the off-diagonal items below the diagonal. An example correlation scatter plot is shown in Fig. 8A. The distribution of *r* value of 1000 simulation is shown in Fig. 8B.

### The link between linear Fisher information and the discrimination thresholds of the optimal linear decoder

Fisher information can be converted to orientation discrimination threshold (Δθ):

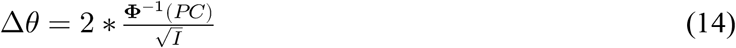

where *I* is Fisher information, *PC* is the percent correct to which the threshold corresponds, and Φ^−1^ is the inverse cumulative normal function. Given 75% accuracy, 1 rad^-2^ information corresponds to the discrimination threshold Δ*θ* of 1.349 radians (i.e., ∼77°).

## AUTHOR CONTRIBUTIONS

R-Y.Z. conceived, designed and performed research; R-Y.Z. wrote the first draft of the paper. R-Y.Z., Zi-Jian Cheng, and W-H. Z. edited the paper.

## ACKNOWLEDGEMENTS

We thank the team of the Studyforrest project (http://studyforrest.org/) to acquire and share the fMRI data. We thank Kendrick Kay, Peter Bandetinni, and Alex Pouget for valuable comments on the manuscript. This work was supported by the National Natural Science Foundation of China (32100901), Shanghai Pujiang Program (21PJ1407800), Natural Science Foundation of Shanghai (21ZR1434700), the Research Project of Shanghai Science and Technology Commission (20dz2260300) and the Fundamental Research Funds for the Central Universities (to R.-Y.Z.).

## SUPPLEMENTAL INFORMATION for

**Figure S1.**
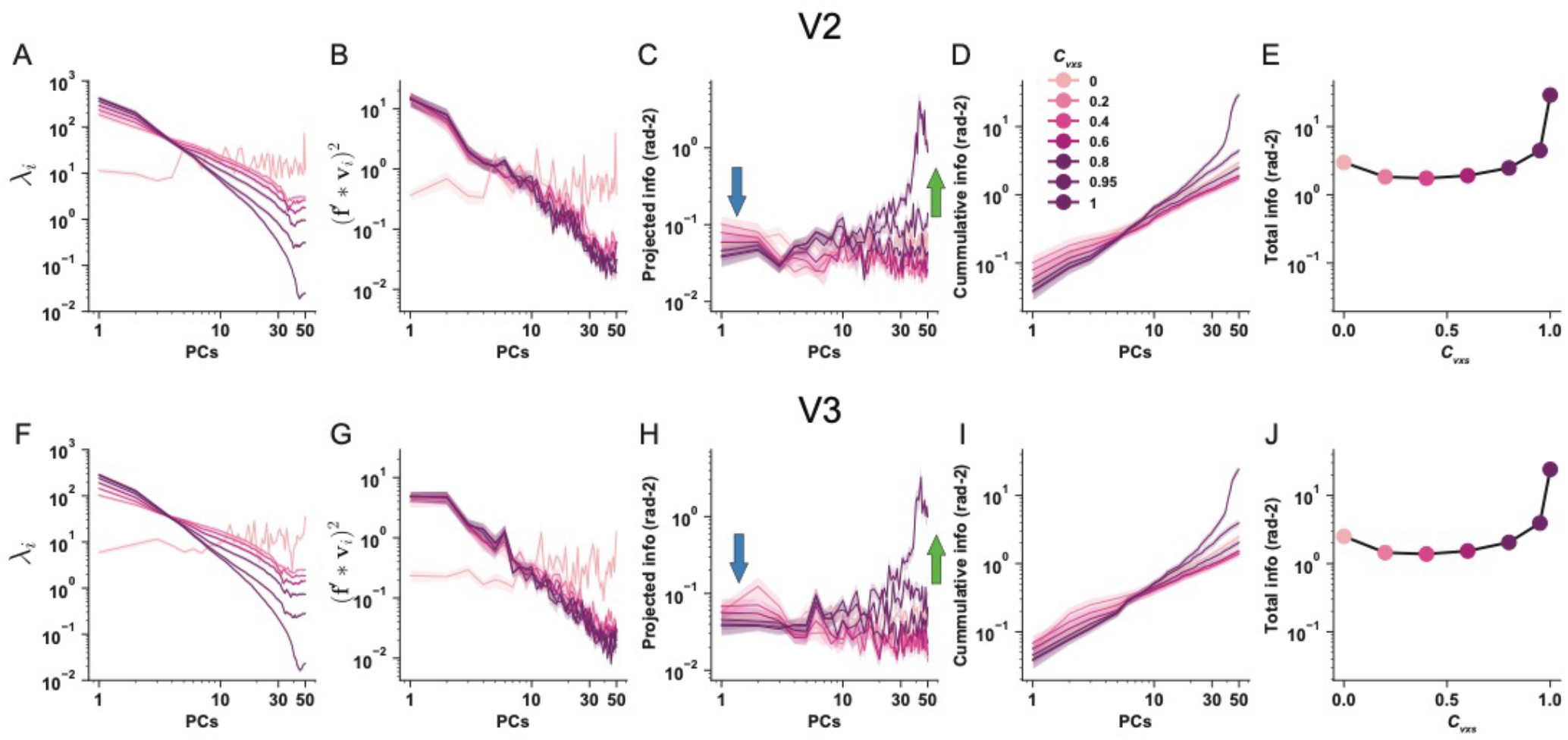
Eigen-decomposition of information in realistic voxel responses in V2 (upper row) and V3 (bottom row). The panel arrangement in each row is similar to Fig. 8K-O in the main text. In both V2 and V3, RCs reduce information on high-variance PCs (indicated by the blue arrows) but disproportionally enhance information on low-variance PCs (indicated by the green arrows), leading to a total higher information. For each line in all panels, we calculated 72 independent samples (6 subjects x 2 hemispheres x 6 pairwise comparisons). There are 6 pairwise comparisons because of the four stimulus orientations. The solid lines and the shaded areas represent the mean and the S.E.M of the samples. The error bars are very small and can be barely seen in ***panels E&J***.

**Figure S2.**
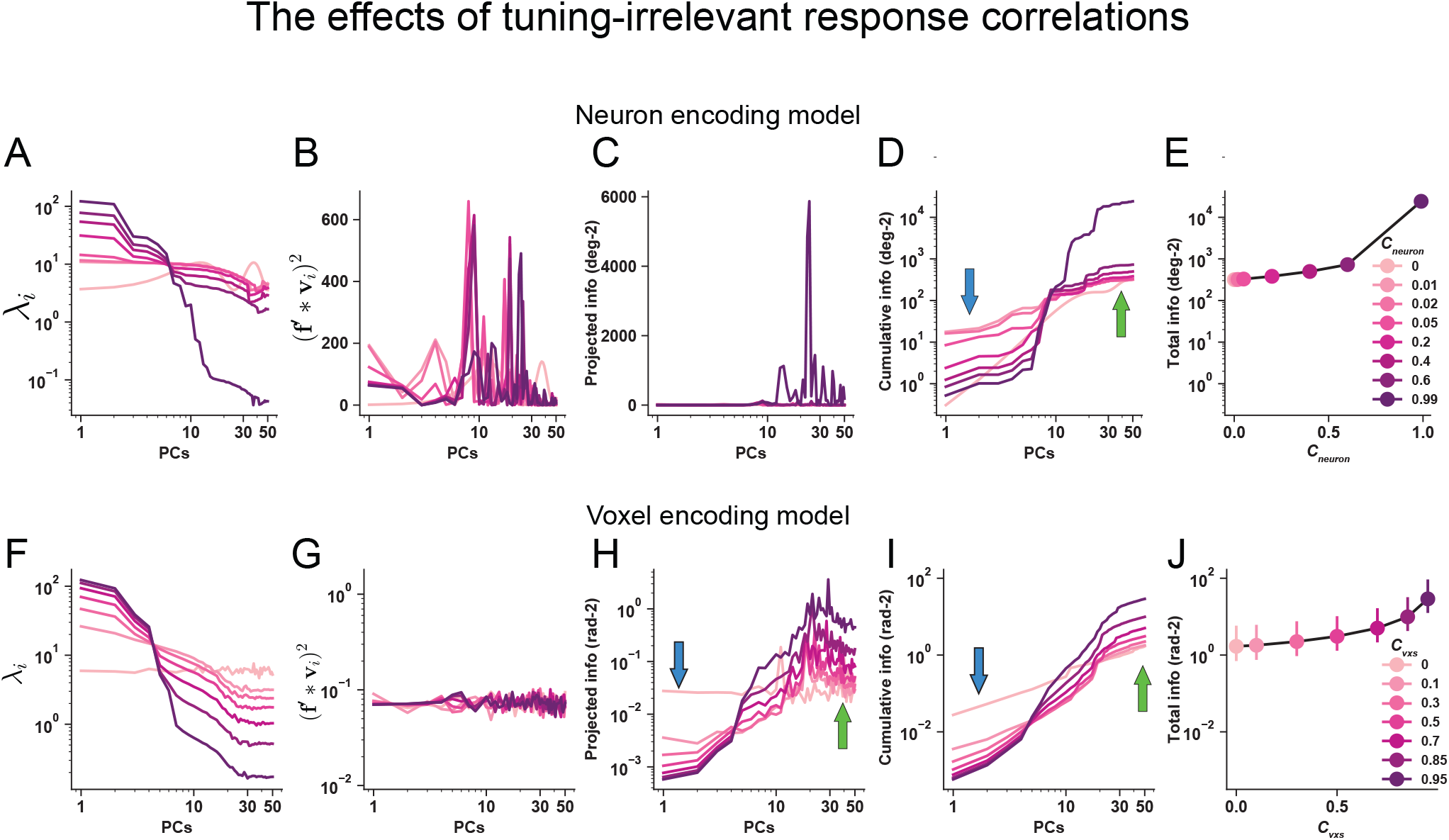
Beneficial effects of tuning-irrelevant response correlations on stimulus information in both neuronal (***Panels A-E***) and voxel populations (***Panels F-J***). The arrangement of the panels is similar to Fig. 7 in the main text. For tuning-irrelevant RCs, we replicate the information reduction on high-variance PCs (indicated by the blue arrows) and information enhancements on low-variance PCs (indicated by green arrows). However, the information reduction is minimal and the information enhancements dominate, resulting in overall beneficial effects of RCs. The error bars in ***Panels J*** represent the 95% confidence intervals of 100 independent simulations.

